# A computational framework with voltage-dependent synaptic function explains LTP-dominant plasticity during functional electrical stimulation therapy

**DOI:** 10.64898/2026.06.15.732206

**Authors:** Michael C. Howard, Kei Masani, Milad Lankarany

## Abstract

Functional Electrical Stimulation (FES) therapy is a widely used neurorehabilitation technique that restores motor function by delivering electrical stimulation to target muscles during voluntary contraction. Despite its clinical effectiveness, the mechanisms by which FES therapy induces neuroplasticity remain poorly understood. Previous work has proposed that positive plasticity arises from Hebbian interactions at corticospinal-motoneuronal synapses when voluntary descending motor commands coincide with antidromic firing elicited by FES therapy. However, if spike-timing-dependent plasticity (STDP) is assumed to underlie this Hebbian mechanism, an unresolved question remains: why does FES therapy produce long-term potentiation (LTP) reliably, rather than the mixture of LTP and LTD predicted from classical STDP rules?

Here, we test the hypothesis that interactions between voluntary descending spikes and stimulation-evoked antidromic spikes generate multi-spike patterns that bias plasticity toward potentiation. To investigate this mechanism, we developed a computational framework implementing a voltage-dependent plasticity rule that incorporates postsynaptic membrane dynamics and higher-order spike interactions. This framework enables simulation of synaptic plasticity during FES therapy while systematically varying stimulation frequency, input heterogeneity, and spike timing structure.

Our simulations show that voltage-dependent dynamics strongly bias synaptic changes toward LTP during FES therapy-like conditions. In particular, physiological interspike interval variability promotes potentiation, whereas highly regular inputs bias synapses toward depression. These results indicate that postsynaptic voltage dynamics and spike-interaction structure, rather than pairwise spike timing alone, govern plasticity outcomes during FES therapy.

Our findings provide a mechanistic explanation for why FES therapy reliably induces LTP-dominant plasticity and offer a computational framework for optimizing neuromodulation therapies.

## 1 Introduction

Functional Electrical Stimulation (FES) therapy is a neurorehabilitation technique in which electrical stimulation is delivered to target muscles during voluntary attempts to perform functional movements. By pairing electrical stimulation with task-related motor commands, FES therapy can produce long-lasting improvements in motor function and corticospinal excitability after neurological injury. Experimental studies have shown that combining voluntary effort with electrical stimulation induces stronger and more persistent plasticity than either of these interventions alone (Barsi et al., 2008; Thompson & Stein, 2004; Yamaguchi et al., 2012). Despite these clinical successes, the mechanisms by which FES therapy induces long-lasting corticospinal plasticity remain poorly understood (Furlan et al., 2022; Popovic et al., 2022; Roman et al., 2023).

One influential hypothesis regarding the mechanism of FES therapy was proposed by Rushton (2003), who suggested that positive neuroplastic changes arise from Hebbian plasticity at corticospinal–motoneuronal synapses (Fig. 1a). In this framework, voluntary descending motor commands provide presynaptic activity, while electrical stimulation evokes antidromic spikes in motor axons that serve as postsynaptic activation. When these signals coincide, synaptic strengthening may occur through Hebbian mechanisms. This hypothesis provides a compelling FES therapy-specific explanation for why combining voluntary effort with electrical stimulation can induce long-lasting plasticity.

**Fig. 1.**
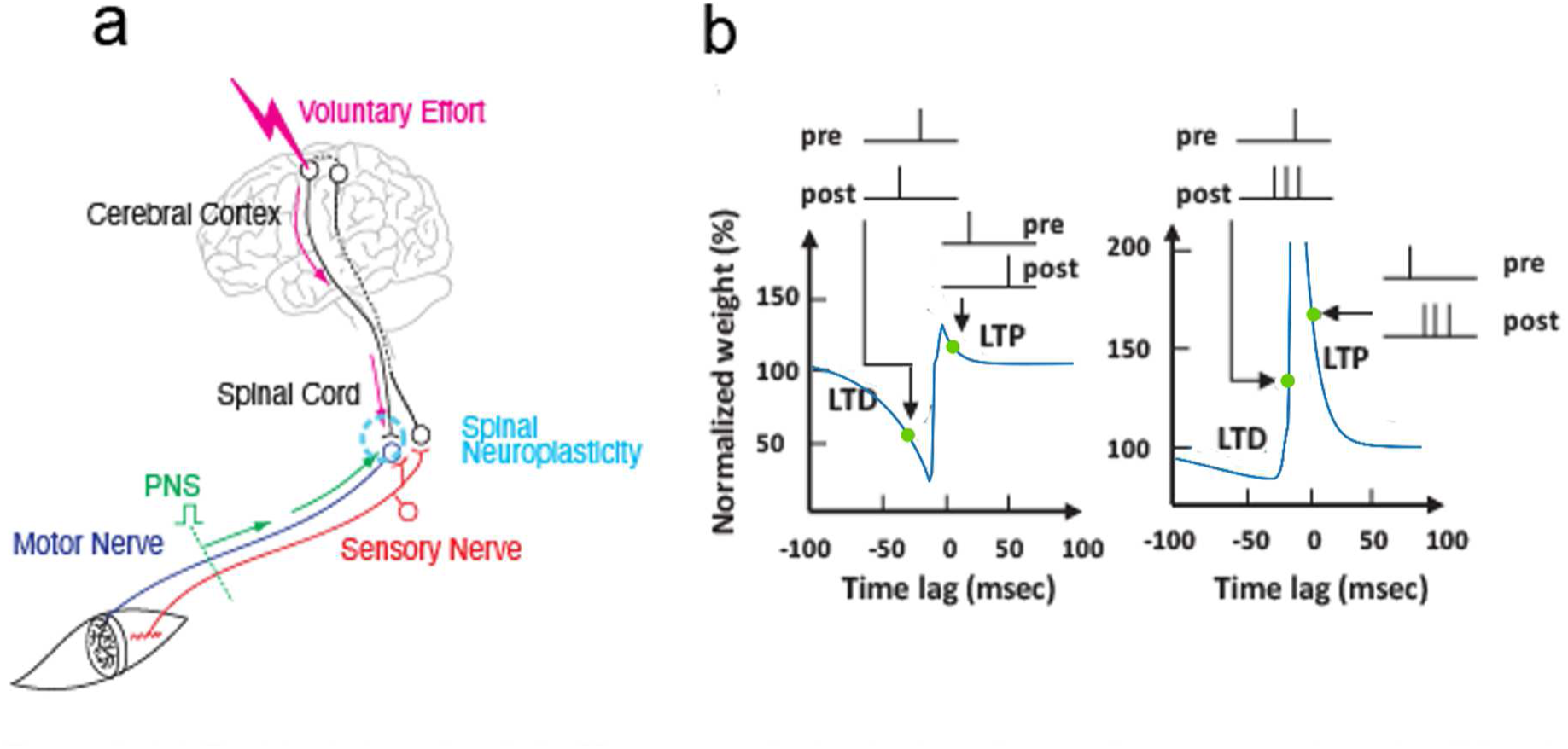
Limits of pair-based STDP in explaining FES plasticity. (a) Rushton’s hypothesis (Rushton, 2003) proposes that voluntary descending commands and antidromic spikes generated by functional electrical stimulation (FES) interact at corticospinal–motoneuronal synapses to induce Hebbian plasticity. (b) Classical spike-timing–dependent plasticity (STDP) predicts that synaptic changes depend on the relative timing of pre- and postsynaptic spikes, producing long-term potentiation (LTP) or long-term depression (LTD) depending on spike order (left). Under more realistic spiking conditions, higher-order spike interactions and postsynaptic voltage dynamics can bias plasticity toward potentiation-dominant outcomes (right). Green markers indicate the magnitude and direction of synaptic weight changes for representative spike pairings redrawn from experimental data shown in Kampa et al. (2006) and Froemke and Dan (2002). Curves were generated using the STDP model of Clopath et al. (2010; see Supplementary Information).

However, when interpreted in the context of spike-timing–dependent plasticity (STDP), which is widely considered a mechanistic implementation of Hebbian plasticity, Rushton’s hypothesis raises an unresolved question. Classical STDP rules predict that synaptic changes depend on the precise timing between presynaptic and postsynaptic spikes, producing either long-term potentiation (LTP) or long-term depression (LTD) depending on spike order (Fig. 1b). Because FES therapy generates complex spike interactions between voluntary descending activity and stimulation-evoked antidromic firing, classical STDP would predict a mixture of LTP and LTD. In contrast, experimental studies of FES therapy consistently report potentiation-dominant plasticity (Carson & Kennedy, 2013; Tolmacheva et al., 2019; Mezes et al., 2020; Furlan et al., 2022). Why FES therapy reliably produces LTP rather than a mixture of LTP and LTD therefore remains an open mechanistic question.

A new approach is therefore needed to investigate the higher-order spiking dynamics that underlie synaptic weight changes in FES therapy. Here we adopt the voltage-dependent plasticity rule proposed by Clopath et al. (2010) as an experimentally grounded alternative to classical Hebbian STDP models (Fig. 1b). Unlike pair-based Hebbian rules, this framework incorporates both presynaptic activity and the dynamics of the postsynaptic membrane potential, enabling it to capture voltage-dependent mechanisms of LTP and LTD (Clopath et al., 2010). This property makes the model particularly suited for FES therapy, where stimulation-evoked antidromic spikes interact with voluntary descending activity to shape postsynaptic depolarization. Using this model, we examine how interactions between voluntary activity and stimulation-evoked spikes influence synaptic plasticity and how factors such as stimulation frequency and input heterogeneity modulate these effects.

## 2 Methods

To investigate the role of synaptic plasticity under functional electrical stimulation (FES), we implemented a spiking neural network model of corticospinal–motoneuronal interactions in NEST Simulator (v3.6.0; Villamar et al., 2023) using Python 3.11.8. All simulations were performed with a temporal resolution of 0.025 ms. Spike timings were rounded to this resolution.

### 2.1 Neuron Model

There were two neuron types used in NEST Simulator: parrot neurons, and adaptive exponential integrate and fire (AdEx) neurons as described in Brette and Gerstner (2005). Parrot neurons simply relay their exact input signal to their targets. The AdEx model was chosen because it captures key electrophysiological features of spinal motor neurons, including spike-frequency adaptation and depolarizing afterpotentials, making it well-suited for voltage-dependent plasticity. Since postsynaptic voltage trajectories critically determine the sign and magnitude of weight changes under the Clopath rule, a voltage-based model was necessary to test FES-induced plasticity. In the AdEx model, the neuron’s membrane potential *u* is governed by an exponential integrate-and-fire equation with adaptation:

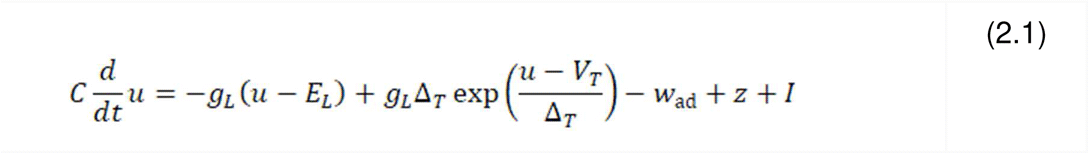

such that *C* is the capacitance, g_L_ is the leak conductance, *E_L_* is the equilibrium potential, *I* is an external stimulating current, and *w_ad_* is an adaptation current, increasing after each spike to contribute a hyperpolarizing effect. The exponential term 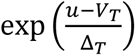 describes the rapid Na channel activation when *u* surpasses an activation threshold *V_T_*, controlled by a slope *Δ_T_*.

The adaptation variable w_ad_ is governed by the equation

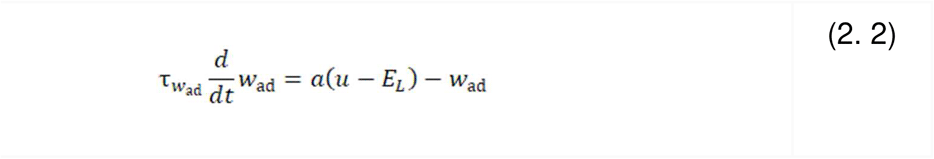

with time constant 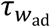 and *a* controlling adaptation strength relative to the membrane potential.

When *u* reaches a peak (e.g., 100 mV above rest), *u* is reset to a lower value *V_T_rest_*, and *w_ad_* is increased by *b* to represent post-spike adaptation. As in Clopath and Gerstner (2010), the AdEx model was extended to include *z*, which represents a depolarizing spike afterpotential (DSAP). At each spike, *z* is set to *I_sp_*, and is modeled with a time constant τ_z._ This DSAP effect, represented by *z*, is relevant in plasticity experiments as it keeps the membrane potential slightly elevated after a spike. This prolonged trace of activity can impact synaptic plasticity mechanisms by maintaining depolarization near the threshold for a brief period, influencing the neuron’s likelihood to spike in response to subsequent inputs. *z* can be understood to represent *I_NaP_*, or a slow non-inactivating Na current (Magistretti & Alonso, 1999).

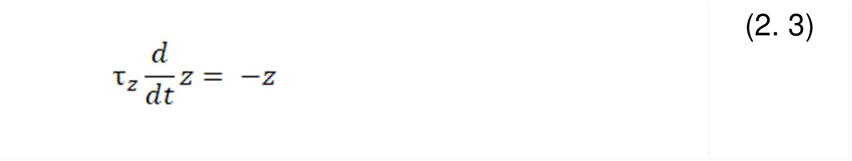

Lastly, the model included an adaptive threshold that jumps to *V_T_max_* after spiking and then decays to *V_T_rest_* with a time constant of *τ_Vt_*.

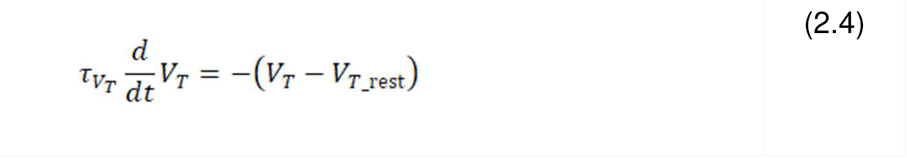

Neuronal parameters used for all experiments are shown in Table 1.

**Table 1:** Parameters of the adaptive exponential integrate-and-fire (AdEx) neuron model used in all simulations. Parameters were taken from Clopath and Gerstner (2010).

| $C$<br>(pF) | $g_L$<br>(nS) | $E_L$<br>(mV) | $\Delta T$<br>(mV) | $V_{T\_rest}$<br>(mV) | $\tau_w$<br>(ms) | $a$<br>(nS) | $b$<br>(pA) | $I_{sp}$<br>(pA) | $\tau_z$<br>(ms) | $\tau_{Vt}$<br>(ms) | $V_{T\_max}$<br>(mV) |
| --- | --- | --- | --- | --- | --- | --- | --- | --- | --- | --- | --- |
| 281 | 30 | -70.6 | 2 | -50.4 | 144 | 4 | 80.5 | 400 | 40 | 50 | -30.4 |

### 2.2 Synaptic Plasticity Model

The plasticity model implemented in this study follows the framework described by Clopath and Gerstner (2010), where synaptic modifications are influenced by the timing of presynaptic spikes and the dynamic, filtered postsynaptic membrane potential. The model separately handles long-term potentiation (LTP) and long-term depression (LTD), each contributing to adjustments in synaptic weight w, representing the connection strength from presynaptic to postsynaptic neurons. For physiological accuracy, w is constrained within bounds from 0 to a maximum value w_max_.

#### 2.2.1 Depression

Synapse depression occurs if a presynaptic spike arrives when the neuron has been depolarized. Mathematically, the presynaptic spike train *X*(*t*) = ∑*iδ*(*t* - *ti*) consists of delta pulses at spike times t_i_, while the postsynaptic membrane potential u is low-pass filtered with a time constant τ_-_.

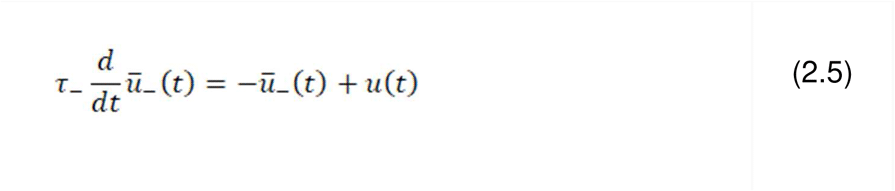

Depression is induced if the postsynaptic trace ū₋ exceeds a threshold θ_−_ at the time of presynaptic spike arrival. This can occur following (1) a recent postsynaptic spike, (2) compound excitatory postsynaptic potentials (EPSPs) from synaptic inputs, or (3) a depolarizing experimental input. The weight change is governed by:

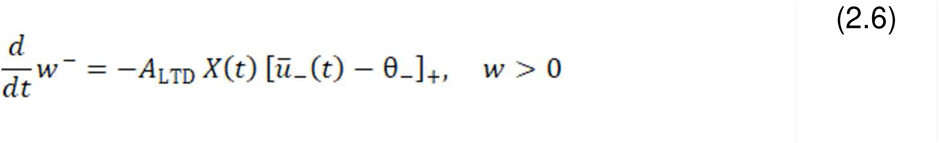

where *A_LTD_* is the depression amplitude. Values in [*x*]_+_ equal *x* when *x* is positive and 0 otherwise. Weight reduction stops when w reaches zero.

#### 2.2.2 Potentiation

Synaptic potentiation occurs if three conditions are met: (1) the momentary postsynaptic voltage *u* exceeds a positive threshold *θ_+_*; (2) ū_+_ (the low pass filtered excitatory postsynaptic potential trace) is greater than a negative threshold, *θ_-_*; and (3) a presynaptic spike occurred recently, leaving a trace represented by x̅. This trace is phenomenological and might reflect factors such as glutamate binding or NMDA receptor upregulation. Potentiation is described by the following equation:

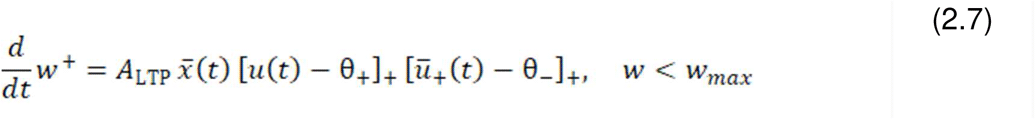

ū_+_ is similar to ū₋ but with a time constant τ_+_. Additionally, x̅ represents a low-pass filtered version of the presynaptic spike train, with a time constant of *τ_x_*.

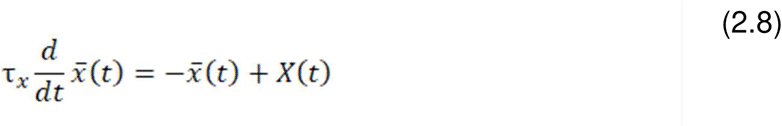

The total equation for weight change is shown below with the hard bounds 0 < w < w_max._

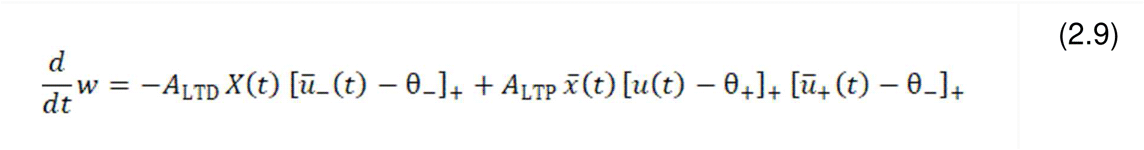

### 2.3 Network implementation and simulation configurations

Simulations were conducted across multiple network configurations to examine plasticity at different levels of organization. In the population configuration, 10 upper motor neurons (UMNs) projected to a population of 500 lower motor neurons (LMNs). Connectivity was defined using fixed indegree sampling such that each LMN received input from five randomly selected UMNs via plastic synapses. Unless otherwise specified, all synapses were initialized with a weight of 20 (arbitrary units). In simulations with heterogeneous initial weights, synaptic weights were instead independently sampled from a normal distribution (μ = 20, σ = 3).

To more specifically investigate plasticity mechanisms, a reduced configuration was used consisting of two UMNs projecting 5 synapses each to a single LMN, thereby isolating synaptic and postsynaptic dynamics at the level of a single postsynaptic neuron.

### 2.4 Presynaptic input generation

Two classes of presynaptic input were used to dissociate the effects of stochastic firing and controlled temporal structure.

First, stochastic inputs were generated using sinusoidally modulated Poisson processes with a mean firing rate of 5 Hz. This formulation captures irregular but temporally structured corticospinal activity and was used in simulations examining frequency-dependent plasticity and network-level robustness.

Second, deterministic spike trains were constructed with fixed interspike intervals (200 ms; 5 Hz baseline rate) and used to precisely control ISI variability. Temporal structure was manipulated by independently jittering each spike time according to:

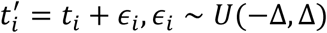

where Δ defines the maximum spike-time deviation. This manipulation preserves the mean firing rate while inducing controlled variability in interspike intervals. Δ was varied up to 100 ms, ensuring that jittered spike times remained within adjacent intervals without overlapping neighboring events.

### 2.5 Synaptic delays

To capture physiological dispersion in corticospinal conduction, synaptic delays were sampled from a normal distribution (μ = 50 ms, σ = 30 ms) in simulations involving stochastic presynaptic input.

### 2.6 FES therapy stimulation protocol

Postsynaptic neurons received stimulation via externally applied current pulses designed to evoke action potentials, modeling antidromic activation of motor neurons during FES. This protocol captures the core FES therapy mechanism whereby voluntary descending activity interacts with stimulation-evoked spikes to drive synaptic plasticity.

Stimulation was delivered as periodic current injections at frequencies ranging from 5 to 200 Hz. Pulse amplitudes were set to reliably elicit postsynaptic spikes. Stimulation was applied for 50 s (20 s – 70 s) within an 80 s simulation window.

### 2.7 Inhibitory input

Inhibitory tone was modeled as a Poisson spike train projecting to LMNs with fixed synaptic weight (−5) and 1 ms delay. The rate of inhibitory input was scaled as:

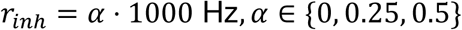

such that increasing a increased the frequency of inhibitory synaptic events while preserving their amplitude.

### 2.8 Convergent input heterogeneity

To examine how interactions between multiple inputs shape synaptic plasticity, simulations included configurations with two independent presynaptic inputs projecting to a single LMN. Each input was assigned an independent jitter parameter (Δ_l_, Δ_2_), allowing systematic manipulation of heterogeneity across converging pathways.

This framework enabled testing of how differences in temporal structure between inputs influence plasticity outcomes under identical stimulation conditions.

### 2.9 Synaptic parameters

Parameters for the voltage-dependent plasticity model were taken directly from Clopath and Gerstner (2010) and were not tuned in the present study (Table 2).

**Table 2.** Parameters of the voltage-dependent STDP rule. Values were taken directly from the original model and control the amplitudes and time constants of potentiation (LTP) and depression (LTD) processes. Parameters were taken from Clopath and Gerstner (2010).

| $\theta_-$<br>(mV) | $\theta_+$<br>(mV) | $A_{LTD}$<br>(1/mV) | $A_{LTP}$<br>(1/mV<sup>2</sup>) | $\tau_X$<br>(ms) | $\tau_-$<br>(ms) | $\tau_+$<br>(ms) |
| --- | --- | --- | --- | --- | --- | --- |
| -70.6 | -45.3 | $21e^{-5}$ | $65e^{-6}$ | 13.3 | 13.8 | 58.7 |

## 3 Results

### 3.1 FES therapy-like stimulation produces frequency-dependent synaptic plasticity

Figure 2 illustrates the synaptic plasticity that emerges under FES therapy-like stimulation conditions. Voluntary descending activity and stimulation-evoked antidromic spikes converged at corticospinal–motoneuronal synapses, allowing interactions between endogenous motor commands and FES inputs.

**Fig. 2.**
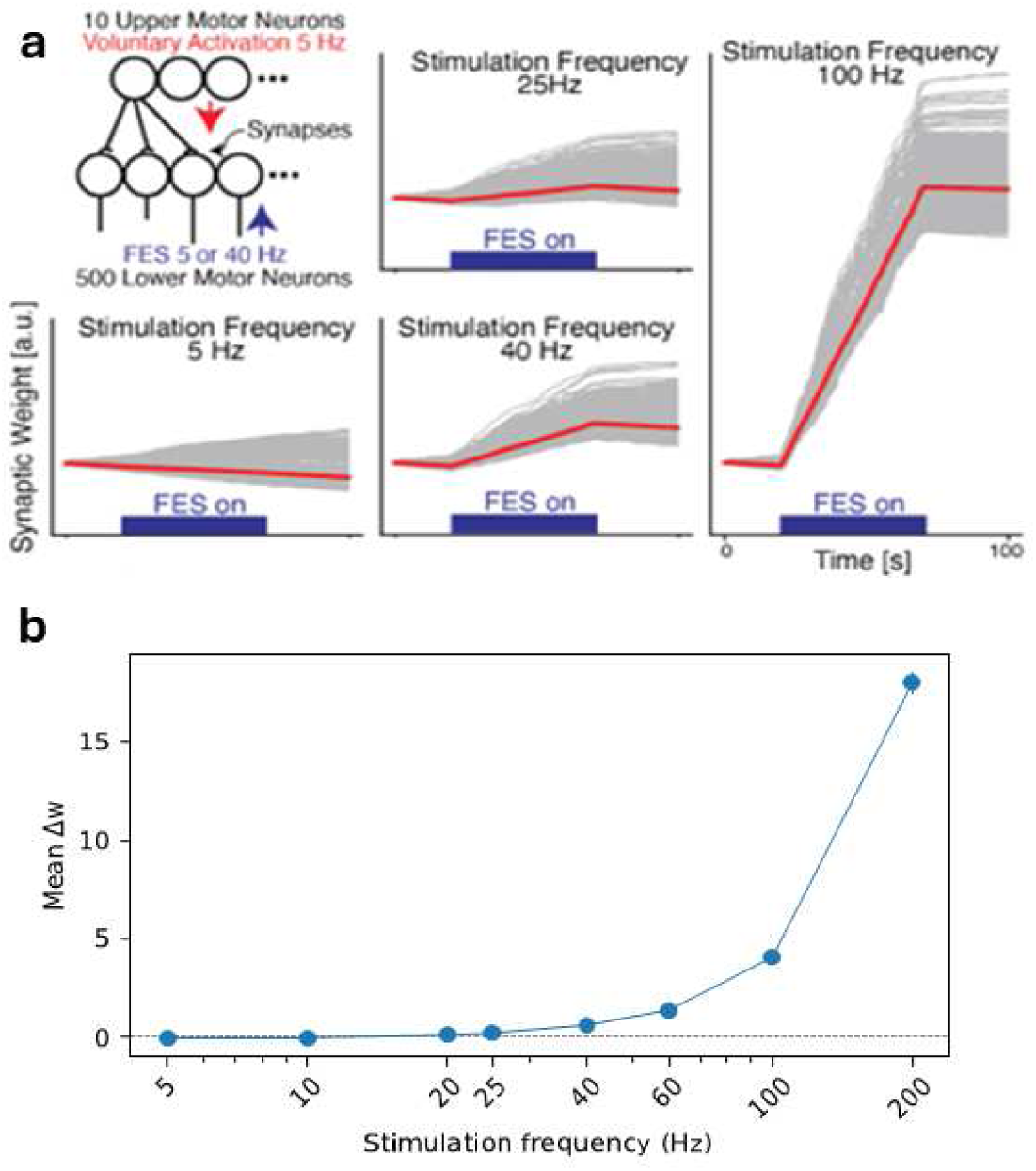
FES Therapy Model. (a) Neural network used in the computational simulation (Left Top), and change of synaptic weights after FES therapy with stimulation intensity of 5, 25, 40, and 100 Hz (other plots). Gray lines are for individual synapses, and the red lines are for the ensemble averages. (b) Mean synaptic weight change (Δw = w_final − w_initial) as a function of stimulation frequency (5–200 Hz). Error bars indicate SEM. Under the voltage-dependent rule, potentiation increases monotonically with frequency, demonstrating that antidromic frequency acts as a control parameter governing the magnitude of plasticity in FES

Representative simulations revealed that synaptic weights evolved differently depending on stimulation frequency (Fig. 2a). At low stimulation frequencies, synaptic changes were small and fluctuated around baseline values. In contrast, intermediate to high stimulation frequencies produced consistent increases in synaptic weights across synapses, indicating potentiation-dominant plasticity.

Population-level analysis further demonstrated a systematic relationship between stimulation frequency and synaptic plasticity (Fig. 2B). Low frequencies produced minimal or mixed plasticity, whereas intermediate frequencies generated robust long-term potentiation across the population. These results indicate that FES therapy-like stimulation biases synaptic plasticity toward potentiation within specific stimulation regimes.

### 3.2 Postsynaptic voltage dynamics, not spike timing, govern frequency-dependent potentiation

Figure 3 examines the mechanisms underlying frequency-dependent plasticity using a reduced configuration consisting of two presynaptic inputs projecting to a single postsynaptic neuron, allowing direct analysis of spike timing, postsynaptic voltage, and plasticity events.

**Fig. 3.**
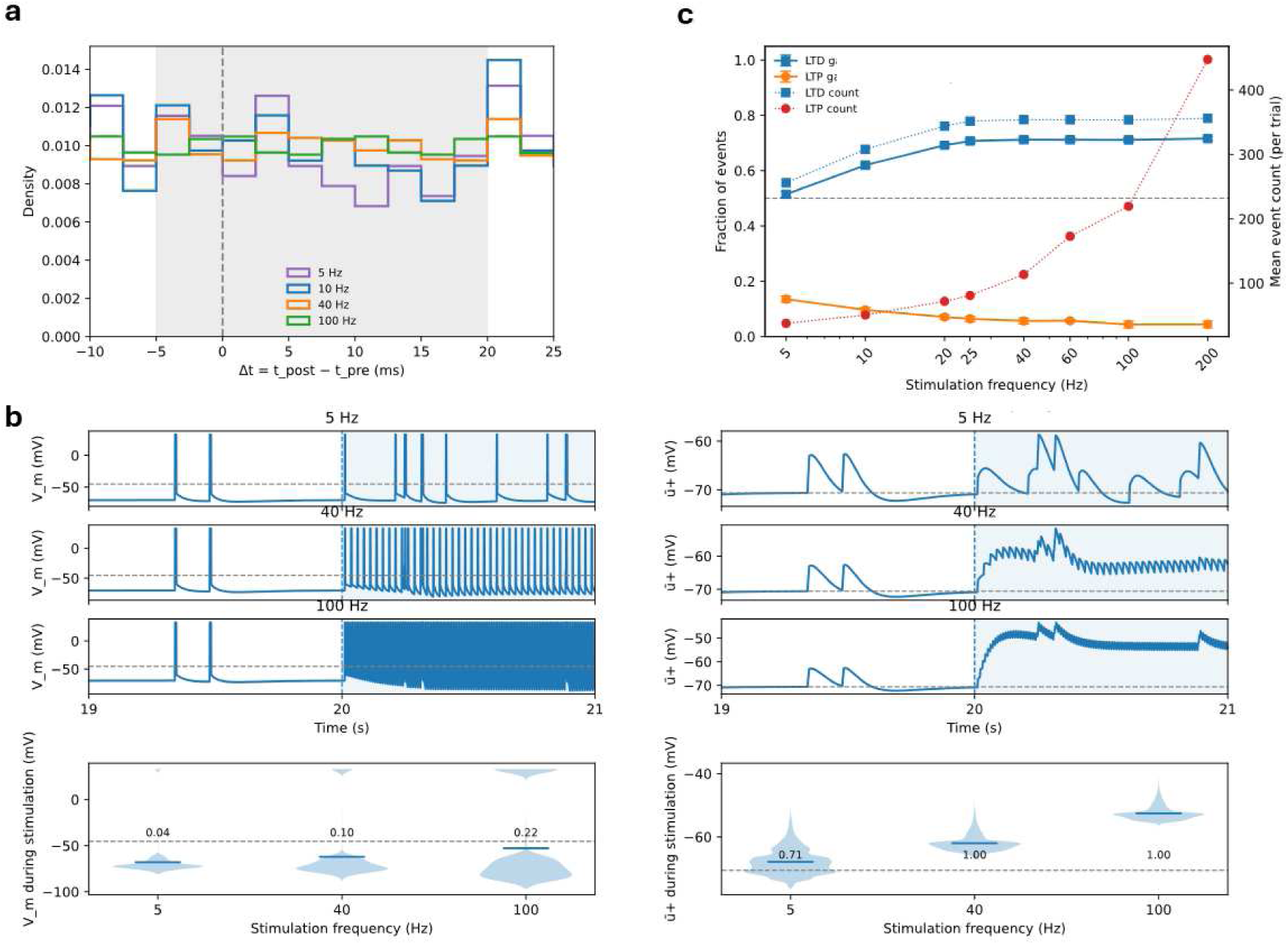
Voltage Dynamics in FES Therapy. (a) Distributions of pre–post spike timing intervals (Δt = t_post − t_pre) across stimulation frequencies. The predicted LTP-favorable window (−5 to +20 ms, shaded) contains a larger proportion of events at lower frequencies, a pattern that would predict stronger potentiation under pair- or triplet-based STDP rules alone, yet these conditions produce weaker or absent LTP. (b) Postsynaptic voltage dynamics during stimulation. Membrane potential traces (V_m) and filtered voltage (ū+) before and after stimulation onset (vertical line, 20 s) show progressively stronger temporal summation with increasing frequency, producing sustained depolarization plateaus. Violin plots of V_m and ū+ during the stimulation window demonstrate that the fraction of time spent above the potentiation thresholds (*θ+*, *θ −* respectively, horizontal dashed line) increases sharply from 5 to 25–40 Hz, indicating enhanced voltage gating of plasticity. (c) Frequency dependence of plasticity events. The fraction of LTP- and LTD-gated events and the absolute count of LTP and LTD events per simulation reveal that, despite saturation of the LTP fraction at 25 Hz, the number of LTP-eligible events continues to rise with frequency due to denser spike interactions.

Distributions of pre–post spike timing intervals (Δt = t_post − t_pre) varied systematically with stimulation frequency (Fig. 3a). Lower stimulation frequencies produced a larger proportion of spike pairings within the canonical LTP-favorable window (−5 to +20 ms), whereas higher frequencies resulted in broader timing distributions with fewer pairings in this range. Despite this, low-frequency stimulation produced weak or mixed synaptic changes, while higher frequencies produced stronger potentiation.

Postsynaptic voltage dynamics showed a distinct dependence on stimulation frequency (Fig. 3b). Increasing stimulation frequency led to greater temporal summation of membrane potential, resulting in sustained depolarization during the stimulation window. Both the membrane potential (V_m) and its filtered component (ū_+_) spent increasing amounts of time above potentiation thresholds as frequency increased from 5 Hz to 25–40 Hz, after which this effect saturated.

This pattern was reflected in the frequency dependence of plasticity events (Fig. 3c). While the fraction of LTP-gated events plateaued at intermediate frequencies, the total number of potentiation events increased with stimulation frequency due to denser spike interactions. As a result, net synaptic weight change increased monotonically with frequency.

Together, these results show that frequency-dependent potentiation tracks changes in postsynaptic depolarization rather than the distribution of pre–post spike timing intervals.

### 3.3 Frequency-dependent potentiation is robust to synaptic heterogeneity and inhibitory tone

Figure 4 tests whether the frequency-dependent potentiation observed in the model persists under more physiologically realistic synaptic heterogeneity and inhibitory tone.

**Fig. 4.**
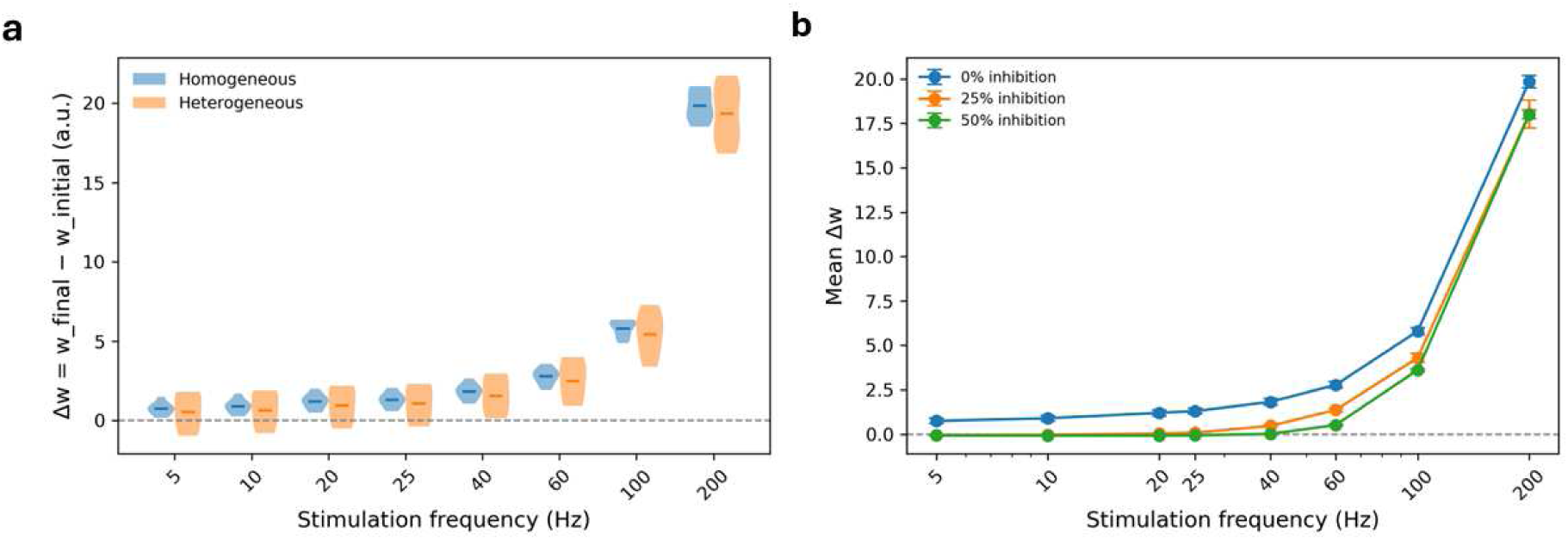
Robustness of Frequency-dependent Potentiation to Synaptic Heterogeneity and Inhibition. (a) Violin plots of synaptic weight change (Δw = w_final − w_initial) across stimulation frequencies for homogeneous (blue) and heterogeneous (orange) initial synaptic weights. Heterogeneity broadens the distribution of outcomes but preserves the monotonic increase in potentiation with frequency. Dashed line indicates no net change. (b) Mean synaptic weight change as a function of stimulation frequency for increasing levels of inhibitory tone (0%, blue; 25%, orange; 50%, green). Inhibition reduces overall potentiation and shifts the response toward higher frequencies but does not alter the ordering of conditions. Across both manipulations, synaptic strengthening increases progressively with frequency, consistent with voltage-dependent gating of plasticity.

Introducing heterogeneity in initial synaptic weights broadened the distribution of synaptic outcomes but did not alter the overall frequency dependence of plasticity (Fig. 4a). Simulations in which synaptic weights were randomly drawn from a normal distribution (μ = 20, σ = 3) produced a wider range of weight changes compared with the homogeneous baseline condition (w₀ = 20 for all synapses). Despite this variability, the mean synaptic response remained unchanged, and potentiation increased monotonically with stimulation frequency.

We next examined the influence of inhibitory tone on plasticity. Adding background inhibitory input to lower motor neurons reduced overall depolarization and attenuated synaptic potentiation across stimulation frequencies (Fig. 4b). Increasing inhibitory input effectively shifted the frequency–plasticity relationship toward higher frequencies, such that stronger stimulation was required to produce comparable potentiation.

Importantly, inhibition scaled the magnitude of plasticity without altering the ordering of stimulation conditions. Across all inhibitory levels tested, higher stimulation frequencies consistently produced larger increases in synaptic weight.

Together, these results demonstrate that frequency-dependent potentiation is robust to physiologically plausible levels of synaptic heterogeneity and spinal inhibition.

### 3.4 Interspike interval variability biases synaptic plasticity toward potentiation

Figure 5 examines how variability in presynaptic spike timing influences plasticity outcomes under FES therapy-like stimulation.

**Fig. 5.**
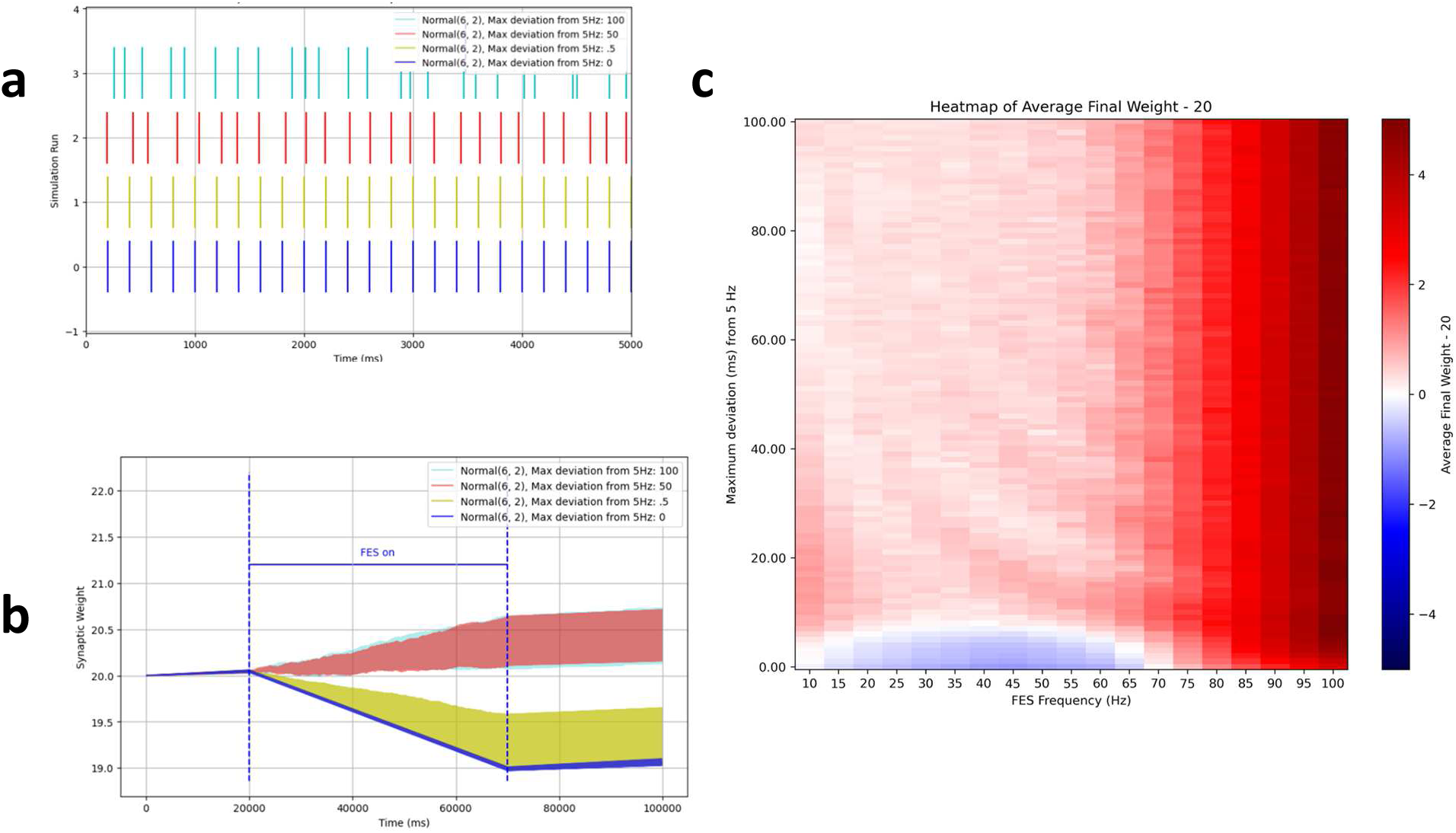
FES Model Performance with Different Input Heterogeneities. (a) Four Poisson spike trains are shown, each with a base frequency of 5 Hz (blue), corresponding to spike times at 200 ms, 400 ms, 600 ms, etc. Spike times deviate by up to half the period (100 ms). Greater deviations from 5 Hz increase variability in interspike intervals (ISI). Color coding represents the maximum deviation: 0 ms (blue), 0.5 ms (yellow), 50 ms (red), and 100 ms (cyan). (b) The FES model simulation is run using each of the spike trains from Panel A as input. The synaptic weight is plotted over time for each simulation. (c) The final synaptic weight is recorded at increasing deviations from 5 Hz and increasing frequency of FES, with darker reds indicating greater potentiation and lighter blues indicating depression.

To isolate the effect of input regularity, we manipulated the variability of presynaptic spike trains while keeping the mean firing rate constant. When presynaptic spikes occurred with highly regular timing, synaptic plasticity was reduced and in some cases shifted toward depression (Fig. 5a). In contrast, introducing physiologically realistic variability in interspike intervals (e.g., maximum deviation from 5 Hz of ≥ 15 ms) produced stronger potentiation across stimulation conditions.

This effect arises because irregular spike timing generates a broader distribution of pre–post interactions and postsynaptic voltage fluctuations. These fluctuations increase the probability that the postsynaptic voltage trace exceeds the potentiation threshold in the voltage-dependent plasticity rule, thereby facilitating long-term potentiation.

Across simulations, intermediate levels of temporal variability consistently produced the largest increases in synaptic weight, whereas highly regular spike trains reduced potentiation and occasionally biased plasticity toward LTD (Fig. 5b).

These findings suggest that physiological variability in neural activity may play an important role in shaping plasticity during FES therapy. In particular, the results raise the possibility that more regular stimulation patterns could be exploited to bias synaptic changes toward depression, a mechanism that may be relevant for therapeutic strategies targeting pathological hyperexcitability such as spasticity.

### 3.5 Input heterogeneity across converging pathways enhances potentiation under FES therapy-like stimulation

Figure 6 examines how synaptic plasticity under FES therapy-like stimulation is shaped by the convergence of excitatory inputs with distinct spike-timing heterogeneity, capturing interactions between physiologically heterogeneous motor activity and regular stimulation-like input.

**Fig. 6.**
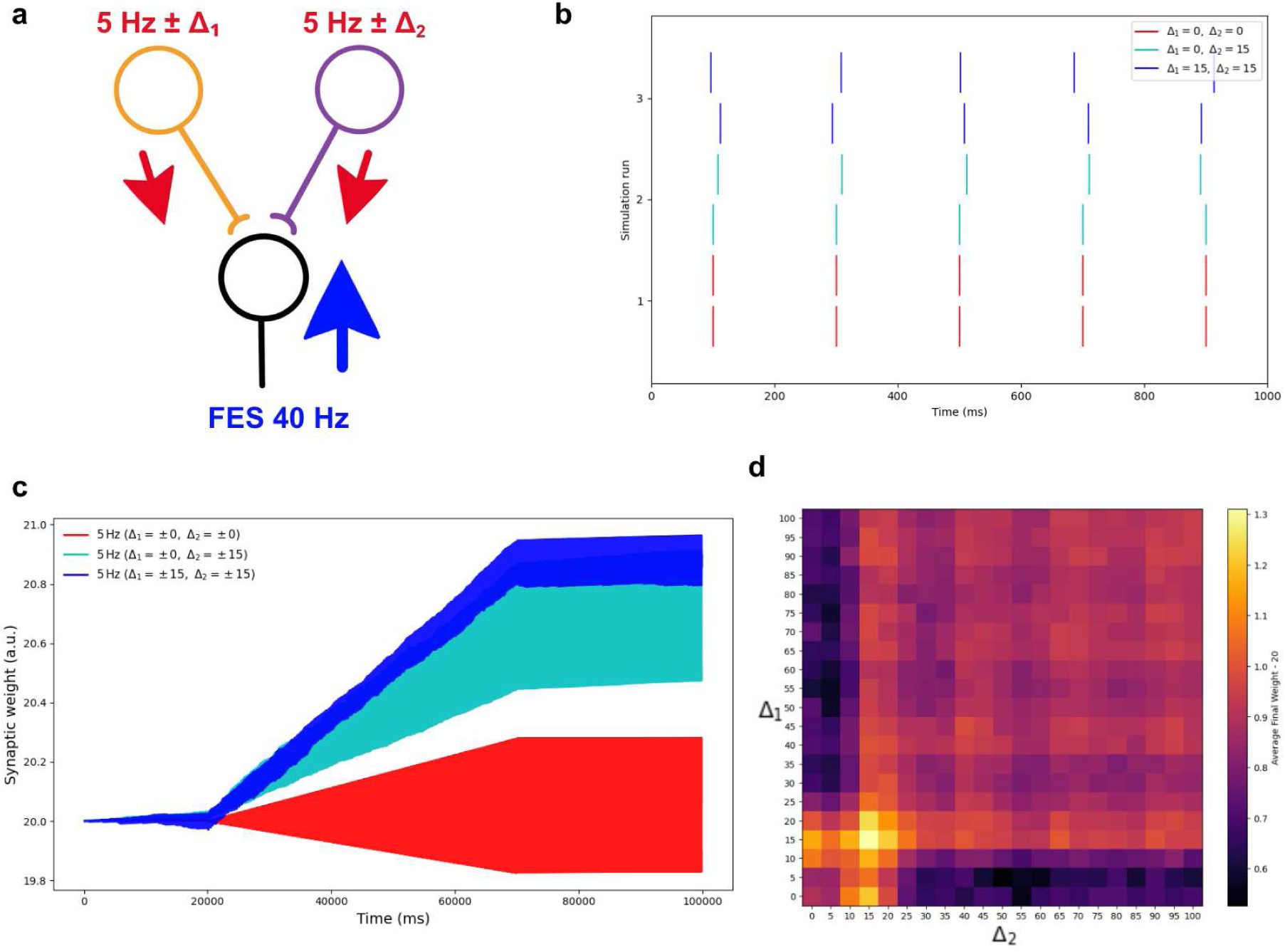
Performance of Two-input FES Therapy Model. (a) Schematic of the experimental setup showing two excitatory inputs converging onto an LMN with antidromic stimulation applied via FES. (b) Example pairs of presynaptic spike trains illustrating different ISI heterogeneity conditions. The first and second inputs are assigned maximum temporal deviations (Δ₁ and Δ₂) from the baseline 5 Hz firing pattern. Example conditions shown are Δ₁ = 0 ms, Δ₂ = 0 ms (red), Δ₁ = 0 ms, Δ₂ = 15 ms (cyan), and Δ₁ = 15 ms, Δ₂ = 15 ms (blue). Increasing values produce greater variability in interspike intervals while maintaining the same average firing rate. (c) Weight changes (Δw) over time for three heterogeneity pairings. (d) Heatmap of final weight changes across heterogeneity pairs under 40 Hz FES. The x- and y-axes represent the heterogeneity assigned to the first and second input, respectively. Warmer colors indicate stronger potentiation.

To isolate the effect of input variability, we simulated two converging presynaptic spike trains with independently controlled ISI jitter while maintaining matched mean firing rates. When one or both inputs exhibited low variability (homogeneous ISIs), potentiation was attenuated and could shift toward depression. In contrast, maximal potentiation occurred when both inputs were physiologically heterogeneous (e.g., maximum deviation ≥ 15 ms), while mixed pairings (0/15 ms) produced intermediate weight changes, indicating that a single low-variability input is sufficient to dampen potentiation.

Two presynaptic parrot neurons each delivered 5 Hz spike trains to a single AdEx-modeled LMN receiving 40 Hz FES (Fig. 6a). Representative spike trains for three heterogeneity pairings (15/15 ms, 0/15 ms, and 0/0 ms) are shown in Fig. 6b, with corresponding weight trajectories in Fig. 6c. A heatmap of final weight change across heterogeneity pairings (Fig. 6d) further shows that potentiation was strongest when both inputs were variable and reduced whenever one input remained low variability.

Together, these results show that under FES therapy-like stimulation, synaptic plasticity depends on the joint temporal structure of converging inputs rather than stimulation alone.

## 4 Discussion

### 4.1 Voltage-dependent plasticity explains LTP-dominant outcomes in FES therapy

FES therapy is widely used in neurorehabilitation, yet its mechanisms remain incompletely understood. A long-standing hypothesis proposed by Rushton (2003) suggests that plasticity during FES arises from spike-timing interactions between voluntary descending activity and antidromic activation of motor neurons. In this framework, descending corticospinal spikes act as presynaptic events while stimulation-evoked antidromic spikes generate postsynaptic firing, creating the conditions for spike-timing-dependent plasticity at corticospinal–motoneuronal synapses (Rushton, 2003). However, classical pair-based STDP rules predict that such interactions should produce a mixture of LTP and LTD, depending on the precise ordering of spikes (Bi & Poo, 1998). This prediction is inconsistent with experimental observations showing predominantly potentiation-like effects following FES therapy.

Our simulations demonstrate that voltage-dependent plasticity provides a mechanistic explanation for this apparent paradox. Using the Clopath plasticity rule (Clopath et al., 2010; Clopath & Gerstner, 2010), the model reproduced both classical STDP curves and higher-order burst-dependent plasticity phenomena observed experimentally (Kampa et al., 2006), validating its suitability for studying stimulation-induced plasticity. When applied to a network model of corticospinal inputs and lower motor neurons, the model predicted that stimulation frequency strongly modulates synaptic weight change, with intermediate and high frequencies producing robust potentiation. These results are consistent with experimental studies showing that combining voluntary effort with peripheral stimulation produces stronger corticospinal potentiation than stimulation alone (Barsi et al., 2008; Thompson & Stein, 2004; Yamaguchi et al., 2012).

Mechanistic analysis revealed that this potentiation is not primarily determined by spike-timing statistics. In our simulations, low stimulation frequencies generated a larger fraction of spike pairs within the classical LTP window yet produced weaker potentiation. Instead, stimulation frequency controlled the postsynaptic voltage state through temporal summation of synaptic and antidromic inputs. Higher frequencies produced sustained depolarization of the membrane potential and its filtered trace, increasing the fraction of time that the voltage exceeded the potentiation threshold of the plasticity rule. As a result, synaptic updates were biased toward LTP despite less favorable spike-pair timing. These findings indicate that postsynaptic voltage dynamics, rather than spike pairing alone, are the primary determinant of plasticity direction during FES.

This voltage-gated mechanism also provides a plausible explanation for the consistent potentiation observed clinically during FES therapy. In the presence of ongoing voluntary activity, stimulation-evoked spikes interact with endogenous motor commands to produce sustained depolarization states in motor neurons. These depolarized states increase the probability that synaptic updates occur under potentiation-permissive conditions, thereby biasing plasticity toward strengthening of corticospinal synapses. Rather than contradicting the Rushton hypothesis, our results suggest that the proposed interaction between voluntary activity and stimulation is indeed critical, but that its plasticity consequences are governed primarily by postsynaptic voltage dynamics rather than pairwise spike timing alone.

Together, these findings provide a mechanistic framework for understanding how stimulation parameters shape plasticity during FES therapy. By incorporating voltage-dependent gating and higher-order spike interactions, the model reconciles the apparent discrepancy between classical STDP predictions and experimentally observed potentiation, and offers a computational approach for investigating stimulation protocols in neuromodulation therapies.

### 4.2 Variability in neural input and circuit conditions shapes stimulation outcomes

In addition to stimulation frequency, our simulations highlight the importance of variability in neural inputs and circuit conditions in determining plasticity outcomes. Although the frequency-dependent potentiation predicted by the model was robust across parameter variations, several additional factors influenced the magnitude and direction of synaptic changes.

First, introducing heterogeneity in synaptic strength broadened the distribution of plasticity outcomes without altering the overall frequency dependence of the system. Likewise, adding inhibitory background input reduced overall depolarization and scaled down potentiation across stimulation frequencies. These findings are consistent with the well-established role of inhibition in regulating spinal excitability and shaping motor neuron responses (Hultborn et al., 2003; Williams & Baker, 2009). Importantly, although inhibition attenuated plasticity magnitude, it did not alter the relative ordering of stimulation conditions, suggesting that stimulation frequency remains a dominant control parameter even under physiologically realistic circuit variability.

Our simulations further revealed that variability in descending input timing strongly influences plasticity outcomes. Motor cortical activity is inherently irregular, with substantial variability in interspike intervals during voluntary movement (Churchland & Shenoy, 2007). When this variability was incorporated into the model, synaptic potentiation increased relative to highly regular spike trains, indicating that physiological heterogeneity promotes potentiation. In contrast, highly regular input patterns biased the system toward weaker potentiation or even synaptic depression. This finding is consistent with theoretical and experimental work suggesting that neural heterogeneity supports robust network dynamics and learning by broadening the range of synaptic interactions (Gast et al., 2024; Perez-Nieves et al., 2021).

These results provide a potential explanation for the variability often observed in clinical and experimental studies of neuromodulation. Differences in cortical firing variability, spinal inhibition, or network heterogeneity across individuals could shift the balance between potentiation and depression during stimulation. Notably, the model predicts that sufficiently high stimulation frequencies can overcome this variability by maintaining sustained postsynaptic depolarization, thereby biasing plasticity toward potentiation regardless of input structure (Fig 5c).

Finally, extending the model to include convergent excitatory inputs demonstrated that interactions between multiple sources of activity can further shape plasticity outcomes. When one input was highly regular, potentiation was attenuated even in the presence of variable endogenous activity. This result parallels paired associative stimulation (PAS) paradigms, in which externally timed stimulation interacts with endogenous neural activity to modulate corticospinal excitability (Alder et al., 2019; Stein et al., 2015).

Together, these findings suggest that stimulation outcomes are determined not only by stimulation parameters, but also by the statistical structure of ongoing neural activity and the state of the underlying circuitry.

Beyond the effects of variability itself, the two-input simulations highlight the importance of interactions between converging sources of activity during FES therapy. Plasticity was influenced not only by the stimulation pattern, but also by the temporal structure of concurrent inputs. This observation is consistent with the original concept proposed by Rushton (2003), in which stimulation-induced activity interacts with ongoing voluntary drive. Our results suggest that the consequences of this interaction are governed by postsynaptic voltage dynamics shaped by the temporal structure of converging inputs, rather than by spike timing alone.

### 4.3 Limitations

Several limitations of the present model should be acknowledged. The simulations focused on simplified corticospinal–motoneuronal interactions rather than full spinal network dynamics. Although background inhibition was included, recurrent inhibitory circuits such as Renshaw cells were not explicitly modeled (Hultborn et al., 2003). In addition, neuromodulatory influences known to affect plasticity thresholds were not incorporated (Pedrosa & Clopath, 2017). Finally, the model does not represent downstream muscular dynamics such as motor unit recruitment, fatigue, or EMG activity, which may influence functional outcomes during FES therapy (Houston et al., 2020; Tian et al., 2024). These simplifications reflect the goal of the study—to isolate the mechanisms governing synaptic plasticity during stimulation—rather than to reproduce the full physiological complexity of the spinal system.

### 4.4 Conclusion

This study presents a voltage-dependent computational framework for investigating synaptic plasticity during functional electrical stimulation (FES). Using the Clopath plasticity rule, the model reproduces the potentiation commonly observed when stimulation is combined with voluntary activity and explains this effect through postsynaptic voltage dynamics rather than spike timing alone.

Our results show that stimulation frequency shapes plasticity by controlling the depolarization state of the postsynaptic neuron, thereby biasing synaptic updates toward potentiation. The model further suggests that variability in neural inputs and circuit conditions can influence stimulation outcomes.

Together, these findings provide a mechanistic framework for understanding FES-induced plasticity and offer a computational approach for exploring stimulation protocols in neuromodulation therapies.

## Acknowledgements

The authors thank members of the Motion & Adaptation Science Laboratory and the Neural Systems & Brain Signal Processing Lab for their constructive feedback during model development and manuscript preparation. M.H. acknowledges the Department of Physiology at the University of Toronto for research support and the Krembil Computational Neuroscience Laboratory (KCN) at the Krembil Research Institute for providing computational resources used in this work. M.H. also thanks David Crompton for training and guidance in computational modeling, and Ryan Ballout for assistance with copyediting the manuscript.

## Author Contributions

Conceptualization: MH, KM, ML. Methodology and Software: MH. Validation and Supervision: KM, ML. Writing — Original Draft: MH. Writing — Review & Editing: ML, KM, MH.

## Data Availability

No datasets were generated or analyzed during the current study.

## Statements and Declarations

## Ethics Approval

Not applicable. This study used only simulated data and did not involve human participants or animals.

## Code Availability

The Python and NEST simulation scripts used to generate the results of this study are available from the corresponding author upon reasonable request. The authors plan to make the full repository publicly accessible following acceptance or upon publication.

