## Supplemental Methods for "A computational framework with voltage-dependent synaptic function explains LTP-dominant plasticity during functional electrical stimulation therapy"

**Supplementary Methods:**

*STDP simulations:*

Pair-based and burst STDP curves (Fig. 1b–c) were generated using the voltage-based plasticity rule of Clopath and Gerstner (2010) implemented in NEST. A single presynaptic neuron was connected to an adaptive exponential integrate-and-fire postsynaptic neuron with Clopath synapses. Simulations used a temporal resolution of 0.1 ms.

*Pair-based protocol:*

Pre–post spike pairs (n = 60) were delivered at fixed inter-burst intervals (5000 ms) following a stabilization period (1 s). The relative timing (Δt) between presynaptic and postsynaptic spikes was systematically varied (−100 to +100 ms). Postsynaptic spikes were evoked via brief current injections (2 ms pulse). Final synaptic weight after all pairings was used to construct the STDP curve.

*Burst protocol:*

Burst-based plasticity was computed using triplet pairings consisting of three postsynaptic spikes paired with a single presynaptic spike across temporal offsets (−100 to 100 ms).

*Parameters:*

Plasticity parameters for pair-based (Fig. 1b) and burst (Fig. 1c) protocols were taken from Clopath and Gerstner (2010) and are summarized in Supplementary Table 1.

Simulations were performed with a parrot neuron connected to a postsynaptic AdEx neuron, with neuronal parameters matching those reported in Table 1.

| | $\theta_-$ (mV) | $\theta_+$ (mV) | $A_{LTD}$ (1/mV) | $A_{LTP}$ (1/mV$^2$) | $\tau_x$ (ms) | $\tau_-$ (ms) | $\tau_+$ (ms) |
|---|---|---|---|---|---|---|---|
| Figure 1b | **−71.3** | **−62.7** | **27e−5** | **12e−5** | **9.6** | **10.5** | **200** |
| Figure 1c | -70.6 | -**65** | **48e$^{-5}$** | **6e$^{-5}$** | **11** | **95** | **5** |

**Supplementary Table 1. Plasticity Parameters.** Parameters were taken from Clopath and Gerstner (2010). Parameter values were fitted to experimental plasticity data in the original study. Parameters that differ from those used in the main simulations (Table 2) are indicated in bold.
